# Theory of Visual Attention (TVA) in Action: Assessing Premotor Attention in Simultaneous Eye-Hand Movements

**DOI:** 10.1101/2020.01.08.898932

**Authors:** Philipp Kreyenmeier, Heiner Deubel, Nina M. Hanning

## Abstract

Attention shifts that precede goal-directed eye and hand movements are regarded as markers of motor target selection. Whether effectors compete for a single, shared attentional resource during simultaneous eye-hand movements or whether attentional resources can be allocated independently towards multiple target locations is controversially debated. Independent, effector-specific target selection mechanisms underlying parallel allocation of visuospatial attention to saccade and reach targets would predict an increase of the overall attention capacity with the number of active effectors. We test this hypothesis in a modified Theory of Visual Attention (TVA; Bundesen, 1990) paradigm. Participants reported briefly presented letters during eye, hand, or combined eye-hand movement preparation to centrally cued locations. Modeling the data according to TVA allowed us to assess both the overall attention capacity and the deployment of visual attention to individual locations in the visual work space. In two experiments, we show that attention is predominantly allocated to the motor targets – without pronounced competition between effectors. The parallel benefits at eye and hand targets, however, have concomitant costs at non-motor locations, and the overall attention capacity does not increase by the simultaneous recruitment of both effector systems. Moreover, premotor shifts of attention dominate over voluntary deployment of processing resources, yielding severe impairments of voluntary attention allocation. We conclude that attention shifts to multiple effector targets without mutual competition given that sufficient processing resources can be withdrawn from movement-irrelevant locations.

## 1. Introduction

When briefly confronted with multiple visual objects in a display, only a subset of the information can be processed due to limited capacities of the visual system. Attention allows the selection of the currently most relevant information in order to guide our behavior adequately (Carrasco, 2011). Over the past decades, different sources of selection bias have been identified, such as selection history, physical salience, and current goals, all of which are integrated in priority maps and determine the spatial deployment of visual attention (Awh et al., 2012). Additionally, researchers have attempted to incorporate different aspects of visual attention into a single theory of attention. One of the most powerful theories is Bundesen’s (1990) *theory of visual attention* (TVA), which accounts for a broad range of findings from behavioral, neurophysiological, and neuropsychological studies on selective attention (Bundesen & Habekost, 2014). Based on the biased competition principle (Desimone & Duncan, 1995), TVA provides a mathematical description of parallel processing of visual objects which compete for selection into visual short-term memory (vSTM). In TVA, attentional selection is achieved by perceptually categorizing the object and storing the categorization in vSTM (for a detailed conceptual description, see Bundesen, 1990; Bundesen & Habekost, 2014). In multi-element displays, where several objects compete for conscious perception and encoding in vSTM (assumed to be limited to *K* different elements), successful selection is determined by the speed of object processing, referred to as *processing rate:* The objects that are processed first win the race for successful storage in vSTM and become available for action control. According to TVA, this rate (*v*(*x, i*)) is composed of three terms: the sensory strength (*η*(*x*, *i*)) of an object *x* belonging to the category *i,* the perceptual bias (*β_i_*) associated with this category, and the relative attentional weight associated with the object (*w_x_*):

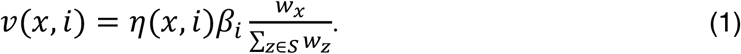

Due to the limited vSTM storage capacity (Luck & Vogel, 1997), it is necessary to prioritize certain objects for further processing. In TVA, prioritization is accomplished by two attentional mechanisms: The *perceptual bias* (*β_i_*) determines how an object is categorized and the assignment of *attentional weights* (*w_x_*) filters which objects are selected for encoding in vSTM (based on the momentary importance of attending to objects belonging to a certain category; Bundesen, 1990; Bundesen & Habekost, 2014). Larger attentional weights thus increase the rate at which an object is selected and its probability to be successfully encoded into vSTM. Because attentional weights are determined for each object in the visual work space, *v*(*x, i*) describes the processing rate of each individual object. The individual processing rates sum up to the overall rate of objects categorization, which is defined as the (attentional) *processing capacity* (Bundesen, 1990; see also **fig. 1b**):

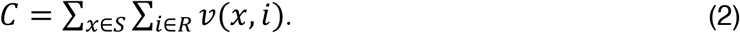

**Fig. 1.**
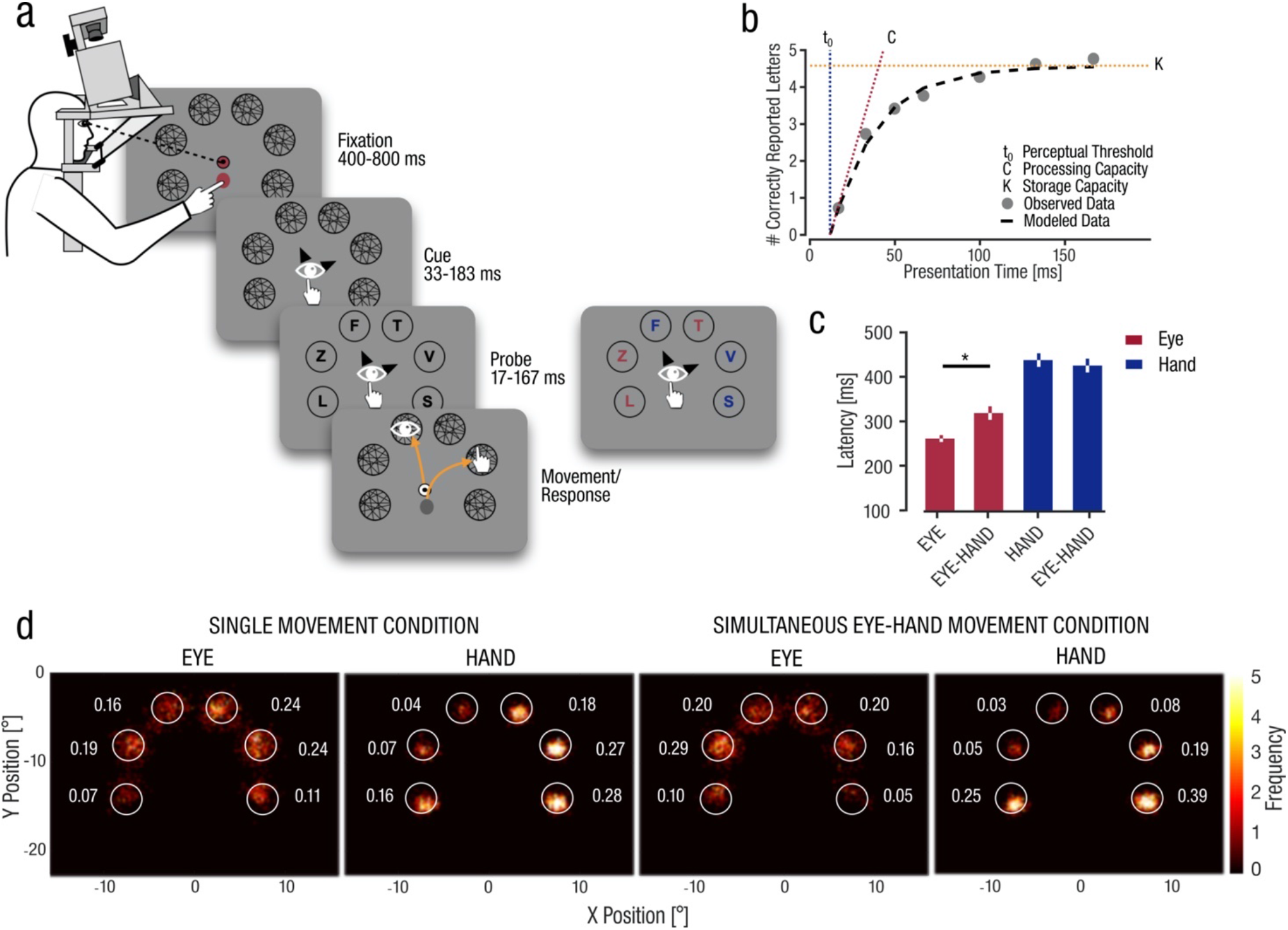
(a) Experimental setup and trial sequence. Participants’ head was fixated on a forehead and chin rest, gaze position of the dominant eye was tracked with an eye-tracker and hand movements of the dominant right hand were recorded by a touch screen. Six circularly arranged masking stimuli appeared. Following a fixation period, two central arrow cues indicated the potential motor targets. Depending on the experimental condition, participants either kept fixating or prepared an eye and/or hand movement toward the indicated target locations. After a random delay, six letters were presented for a varied duration, which participants had to report at the end of the trial. In Experiment 1, letters and masks were black. In Experiment 2, letters and masks were colored in red and blue. (b) Example TVA fitting curve (black line). Grey dots depict the average performance for each presentation time. The intersection of the fitting curve with the x-axis defines the perceptual threshold (*t_0_*), the initial slope at *t_0_* is defined as the processing capacity (*C*), and the asymptote of the curve defines the storage capacity of vSTM (*K).* (c) Average saccade and hand movement latencies across participants. Asterisks represent significant differences (*p* < 0.05). (d) Heatmaps indicate absolute frequencies of eye and hand landing positions across participants and trials. Values represent average proportions across participants for each stimulus location (white circles).

TVA has been applied to determine age-related changes in attention selectivity and capacity across the lifespan (McAvinue et al., 2012), to assess cognitive impairments in various patient groups (Bublak et al., 2005; Duncan et al., 1999; for a review, see Habekost, 2015), and to measure different components of visual attention in healthy populations (Finke et al., 2005). Yet, the TVA-based assessment of visual attention has only been studied under fixation (but see Poth & Schneider, 2018 for TVA-based assessment of transsaccadic competition). Here, we aim to use the theoretical and computational framework of TVA to investigate, for the first time, whether TVA can account for attention mechanisms in relation to selection-for-action (Allport, 1987, Schneider, 1995).

It is well established that both saccades (Deubel & Schneider, 1996; Kowler et al., 1995; Montagnini & Castet, 2007) and hand movements (Baldauf et al., 2006; Deubel et al., 1998; Rolfs et al., 2013) are preceded by obligatory shifts of visual attention towards their motor targets. Some authors even assume that the very purpose of attention is action control (Allport, 1987; Neumann, 1987). Nonetheless, it is unknown how action-coupled attention shifts modify the selection of competing objects for encoding into vSTM. To approach this question, we combined a TVA paradigm with eye-hand movement tasks to investigate how motor preparation affects the deployment of visual attention over multi-element displays. Whereas the classical TVA framework focusses mainly on single-task situations, Logan and Gordon (2001) previously extended TVA to an executive control theory of visual attention (ECTVA) that can account for dual-task situations (for an application of TVA to motor-cognitive dual-task situations, see Künstler et al., 2018). Yet, to the best of our knowledge, our study is the first to evaluate premotor shifts of visual attention via the TVA framework, which offers two key advantages over conventional approaches: First, given the detailed theoretical framework of TVA, we are able to make detailed predictions on how premotor shifts of visual attention affect the selection of competing objects for successful encoding into vSTM. Second, using a multi-element display, TVA allows the assessment of both the overall attention capacity (parameter *C,* captured in the visual processing capacity of letters) and the allocation of attentional resources to each individual object in a display (parameter *v).* Thus, overcoming the limitations of traditional premotor paradigms that can measure attention only at one location at a time (see Hanning et al., 2019 for different approaches), the TVA paradigm allows for a concurrent assessment of premotor visual processing across the entire visual work space.

The advantage of being able to simultaneously evaluate perceptual benefits and costs at motor targets and movement-irrelevant locations becomes particularly evident in the investigation of attentional dynamics during the preparation of combined eye-hand movements: Looking and reaching simultaneously to spatially separate goals requires the selection of multiple movement targets. Previous studies investigating shifts of visual attention – an index of movement target selection – during simultaneous eye-hand movements, observed increased attention at both eye and hand targets (Jonikaitis & Deubel, 2011; Khan et al., 2011; Hanning et al., 2018). Whereas Khan et al. (2011) observed that the eye dominated in guiding attention during simultaneous eye-hand movements and argued in favor of a single, shared attentional resource (see also Nissens & Fiehler, 2018), other studies showed that the attention benefits at two effector targets are not impaired by the necessity to plan simultaneous eye-hand movements versus a single eye or hand movement (Jonikaitis & Deubel, 2011; Hanning et al., 2018). On the one hand, a parallel allocation of attention to multiple effector targets without competition can be explained by separate attentional resources dedicated to each individual effector. In this case, the overall attention capacity should increase with the number of active effectors. On the other hand, the eye and hand target benefits may be accompanied by a withdrawal of attentional resources from non-motor locations – in which case the overall attention capacity would not rise with the number of active effectors.

To unravel this ambiguity, we used TVA to asses visual processing across the entire visual work space, as well as the allocation of attention to the individual motor targets and movement-irrelevant locations. In Experiment 1, we assessed whether the parallel preparation of eye and hand movements to spatially separate goals enhances visual processing at both effector targets simultaneously, and whether such parallel processing benefits would be reflected in increased overall attention capacity. In Experiment 2, we investigated whether attention was withdrawn from non-motor locations in order to allocate processing resources towards the motor targets and whether these costs occurred obligatorily. We used a similar paradigm as in Experiment 1, but in addition to the perceptual and motor task, we gave participants an incentive to voluntarily deploy their attention primarily to a subset of the presented items: different colors (50% red and 50% blue) indicated whether the letters were associated with a high or low monetary reward. This enabled us to investigate the ability to voluntarily deploy attention while preparing eye-hand movement.

Altogether, our experimental design allows us to determine (a) whether attention is deployed to the eye and hand targets in parallel, (b) whether such parallel allocation leads to an increase in the overall attention capacity or is associated with concomitant costs at non-motor locations, and (c) how our goal-directed actions influence our ability to voluntarily attend elsewhere.

## 2. Experiment 1

In this experiment we established the TVA framework as a sensitive tool to measure action-related shifts of visual attention. Specifically, we expected action preparation to increase the attentional weights at the motor targets, reflected in increased processing rates. First, we used this approach to examine if attention is deployed in parallel to eye and hand movement targets and whether the effectors compete for attentional resources. Second, we tested whether the overall attention capacity increases with the number of active effectors.

### 2.1. Methods

#### 2.1.1. Participants

Eight healthy human adults (age range: 24-31 years, four female, one left-handed) participated in the experiment (including authors PK and NMH). All participants had normal or corrected-to-normal visual acuity. We determined our sample size based on previous research on premotor shifts of visuospatial attention using between five and nine participants (e.g., Hanning et al., 2018; Khan et al., 2011). According to previously reported effect sizes (Hanning et al., 2018), a sample size of at least eight is required to detect increased attention deployment at cued saccade (previously reported effect size: *d* = 2.59) and reach (*d* = 1.50) targets in single movement conditions with a power of .95. The experimental protocols were approved by the local ethical review board of the Faculty of Psychology and Education of the Ludwig-Maximilian-University Munich and in accordance with the Declaration of Helsinki. Participants gave written informed consent and were compensated at the rate of €10/hr.

#### 2.1.2. Apparatus

Participants performed the task in a dimly illuminated laboratory, viewing the stimuli binocularly on a 32-inch, 25°-inclined LCD touch display (Elo 3202L Interactive Digital Signage, Elo Touchsystems, Menlo Park, CA, USA) with a refresh rate of 60 Hz and a spatial resolution of 1,920 × 1,080 pixels. The viewing distance to the screen midpoint was 50 cm. An EyeLink 1000 Tower Mount (SR Research, Osgoode, Ontario, Canada) video-based eye tracker recorded the gaze position of participants’ dominant eye at a sampling rate of 1 kHz. To minimize head movements during the task, participants’ heads were stabilized by a combined chin and forehead rest (see **fig. 1a**). The experiment was programmed in Matlab (MathWorks, Natick, MA, USA), using the Psychophysics (version 3.0.14; Kleiner et al., 2007) and EyeLink toolboxes (Cornelissen et al., 2002). Stimulus display and data collection were controlled by a PC (graphics card: NVIDIA GeForce GTX 1070), connected to the eye tracker’s host computer.

#### 2.1.3. Stimuli and Procedure

In order to examine the overall attention capacity and the allocation of attention to motor and non-motor targets, we used a TVA whole report paradigm and measured the categorization performance of letter stimuli presented at six spatially fixed locations while varying the presentation time (17 - 167 ms) between trials. In a randomized block design, participants were instructed at the beginning of each block to either fixate in the center (*FIXATION*), to perform single eye or hand movements (*EYE* or *HAND*), or to perform simultaneous eye and hand movements (*EYE-HAND*). **Figure 1b** depicts the trial sequence. At the beginning of each trial, participants fixated a black-and-white bull’s eye (0.5° visual angle) presented on a uniform grey background (28.2 cd/m^2^) and placed their right index finger on a dark grey oval (0.6° × 0.65°) below the bull’s eye. Six black circles (2° radius), evenly spaced along a 8° virtual arc (± 110° from the vertical) marked the stimulus locations. The trial started once stable eye and finger fixation was detected within a 2.25° radius virtual circle centered on the fixation targets. Six dynamic circular pre-masks composed of randomly oriented black lines were presented at the stimulus locations, which had a random orientation that changed at a rate of 30 Hz. Letter stimuli were embedded within a dynamic mask to prevent visual disruption by the sudden letter on- and offset, which has been shown to interfere with movement preparation (Reingold & Stampe, 2002). Appendix **fig. A1** shows that eye and hand movement latencies were not affected by letter presentation. 400 - 800 ms after pre-mask onset, two black arrow cues (0.25° × 0.5°) were presented around the bull’s eye (with a distance of 1.5°) pointing to two of the six stimulus locations, randomly selected with an equal likelihood for each location. Participants were instructed to ignore these cues in the *FIXATION* condition, as they were not relevant for the task. In the single movement conditions (*EYE* or *HAND*), participants chose one of the two locations and moved the respective effector (eye or hand) to the selected location as fast and as precisely as possible, while the other effector remained at fixation. In the *EYE-HAND* condition, participants moved each effector to one of the cued locations, at free choice. Note that we always presented two cues irrespective of the experimental condition to keep the visual input constant. This way a potential difference in overall attention capacity could not be explained by visual bottom-up processes elicited by the cue. Earlier pilot versions of the task revealed that using different color cues for the different effectors (i.e., to avoid free choice) increased the difficulty of the motor tasks and resulted in high proportions of error trials. Following a stimulus onset asynchrony (SOA) of 33.3 - 183.3 ms after cue onset, the pre-mask was replaced by six black letters (~0 cd/m^2^), selected from a subset of 19 letters (ABEFHJKLMNPRSTVWXYZ). The letters were presented for a duration of either 17, 33, 50, 67, 100, 133, or 167 ms. SOA and letter presentation duration were matched such that the letter presentation always ended 200 ms after cue onset, i.e., within the movement latency. This ensured that the presentation of the letters always occurred shortly before movement onset, when the deployment of attention to movement targets is highest (e.g. Hanning et al., 2018; Montagnini & Castet, 2007). Subsequently, the dynamic masks reappeared and were presented for another 800 ms. Afterwards, participants verbally reported all letters they had perceived and received online feedback if a motor task was not correctly performed.

Participants performed at least 16 experimental blocks (four of each experimental condition) of at least 70 trials each. This resulted in approximately 40 trials per condition and presentation time. The experiment was split into two sessions. We controlled online for broken eye and finger fixation (outside 2.25° from fixation), too short (<170 ms) or too long (>700 ms) movement latencies, and incorrect eye or hand movements (not landing within 2.25° from the indicated targets). Erroneous trials were repeated in random order at the end of each block. If motor performance was incorrect in more than 50% of trials within a block, the whole block was excluded and repeated (this was the case for two blocks for one participant). Overall, participants moved their eyes and hand incorrectly to the same target in 5.2% ± 0.28 (mean ± SE) % of trials, and made other eye movement errors in 13.5 ± 0.78 % (broken fixation: 6.0 ± 0.52 %, incorrect saccade: 7.5 ± 0.37 %) and hand movement errors in 2.0 ± 0.19 % (broken fixation: 0.22 ± 0.03 %, incorrect hand movement: 1.8 ± 0.17 %) of trials.

#### 2.1.4. Data Analysis

Data was analyzed offline using custom-made routines in Matlab (MathWorks, Natick, MA, USA). We measured eye and finger position in x- and y-screen-centered coordinates and determined saccade and reach movement onset, landing time, and position. Saccades were detected based on the eye velocity profiles (Engbert & Mergenthaler, 2006) using a moving average over twenty subsequent eye position samples. Saccade onset and offset were determined when the velocity exceeded or fell below the median of the moving average by 3 SDs for at least 20 ms. Saccade and hand movement latencies were defined in relation to motor cue onset. Offline criteria for correct trials were as follows: (1) eye and hand maintained fixation within a 3° radius centered around the effector’s fixation position until cue onset, and the passive effector(s) in the *EYE*, *HAND*, or *FIXATION* conditions maintained fixation until the end of the trial, (2) eye and hand movements landed no further than 3° from motor targets center and stayed there for at least 50 ms, and (3) no blinks occurred during the entire trial. Based on these criteria, 8.72% of all online accepted trials were discarded.

We modeled the remaining trials according to TVA in two steps. First, we implemented TVA modeling procedures using the libTVA toolbox for Matlab (Dyrholm et al., 2011), in order to measure the allocation of attention to all six individual stimulus locations. To this end, we used the processing rates *v* of each individual object in the visual work space (the rate at which an object is encoded into vSTM; measured in objects per second [Hz]). Perceptual thresholds (*t_0_*) – the longest ineffective exposure duration, below which the probability of reporting any element is zero – were estimated for each participant and experimental condition but were assumed to be constant for the different target types within an experimental condition. The observed and estimated values for uncued locations were averaged to obtain a single measure for the processing rate at uncued, non-motor locations. For the *FIXATION* condition, we also averaged across both cued (not-motor) locations. Second, we assessed the influence of movement preparation on the deployment of visual attention over the whole stimulus display. For this, the following standard TVA parameters were extracted separately for each participant and experimental condition: (1) *t_0_* (perceptual threshold), (2) *K* (visual short-term memory capacity) – the maximum number of items an observer can remember and report, and (3) *C* (processing capacity) – the overall visual processing speed, distributed across all objects in the visual work space according to their attentional weights. Conceptually, the processing capacity is the summed processing rate (*v*) of all items in the visual work space (for exemplary fitting curves and the extracted parameters, see **fig. 1b**). For all analyses, *t_0_* was set to 0 ms if its estimation was < 0 ms.

To evaluate the differences in overall attention parameters across experimental conditions as well as differences in attention deployment within each condition, we ran multiple repeated-measures analyses of variance (rmANOVA). Sphericity was assessed using Mauchly’s test and Greenhouse-Geisser-corrected *p*-values are reported in case of significance. An alpha level of 0.05 was chosen and for post-hoc pairwise comparisons Bonferroni-corrected *p*-values are reported. All statistical analyses were performed in R (version 3.3.2; R Core Team, 2016).

### 2.2. Results

Participants made eye and/or hand movements towards centrally cued targets. Saccades occurred 261 ms (± 22 ms; mean ± STD) after cue onset in the single eye movement (*EYE*) condition – significantly faster as compared to the combined *EYE-HAND* condition (318 ± 43 ms; *t*(7) = −5.41; *p* < 0.001; *d* = 1.91; **fig. 1c**). In general, reach latencies were longer but did not differ between the single (*HAND*) and combined (*EYE-HAND*) movement condition (437 ± 43 ms and 425 ± 44 ms, respectively; *p* = 0.32). Note that whereas previous reports showed highest attention allocation at the motor target shortly before movement onset (Deubel, 2008), we recently showed that attention effects emerged simultaneously after cue onset at both the eye and the hand target (Hanning, et al., 2018). Thus, we do not expect differences in eye and hand movement latencies to affect our results. When analyzing the distribution of saccade and reach endpoints, we observed a clear target selection preference for the two effectors (**fig. 1d**): Participants preferentially chose the upper locations as saccade targets and the lower locations on the right-hand side as reach targets. This differential preference for eye and hand target selection is clearly visible when eye and hand movements were programmed simultaneously (right panels of **fig. 1d**) and might be explained by participants avoiding crossing the two effector paths.

#### 2.2.1. Allocation of Attention at Motor Targets versus Non-Target Locations

The first aim of the current study was to assess how action-coupled shifts of visual attention modify the selection of competing objects for successful encoding into visual short-term memory (vSTM). We expected increased allocation of attention at the motor targets in the single movement conditions (*EYE* and *HAND*). Moreover, based on previous studies finding parallel allocation of attention in simultaneous eye-hand movements (Hanning et al., 2018; Jonikaitis & Deubel, 2011), we expected attention to be equally deployed to the saccade and the reach target in the *EYE-HAND* condition. To keep the visual input constant across experimental conditions and to ensure that differences in attention capacity between conditions cannot be explained by a varying number of visual cues, we always presented two cues in all conditions. We did not expect large attention benefits at the cued locations in the *FIXATION* condition, since the cues were task-irrelevant. To test these hypotheses, we estimated the processing rates (*v*) of items presented at motor targets versus cued and uncued non-targets (for fitted curves of a single subject, see **fig. 2a**). **Figure 2b** shows the extracted *v*-parameters for the different target types in each experimental condition averaged across all participants. For the *FIXATION* condition, a paired t-test between averaged processing rates at cued and uncued locations revealed no significant difference (*t*(7) = −2.17; *p* = 0.067; *d* = 0.77). For the other experimental conditions, we ran separate one-way rmANOVAs with factor *target type*, revealing significant effects in each effector condition between motor targets, cued but not selected locations, and uncued locations (*EYE: F*(2,14) = 29.61; *p* = 0.001; 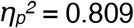, *HAND: F*(2,14) = 47.76; *p* < 0.001; 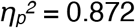, and *EYE-HAND: F*(2,14) = 17.78; *p* < 0.001; 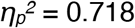). Post-hoc pairwise comparisons indicated a similar deployment of attentional resources in both single movement conditions: processing rates were increased at the motor targets compared to the averaged processing rates at the uncued locations (*t*(7) = −5.52; *p* = 0.003; *d* = 1.95 and *t*(7) = −7.88; *p* < 0.001; *d* = 2.79 for *EYE* and *HAND,* respectively) and at the second cued but not selected location (*EYE* : *t*(7) = −5.40; *p* = 0.003; *d* = 1.91 and *HAND: t*(7) = −6.82; *p* = 0.001; *d* = 2.41). The latter demonstrates that the premotor shifts of attention observed at the motor target cannot be explained by a visual cueing effect. Contrary to the *FIXATION* condition, in both single movement conditions we furthermore observed significant differences between the cued (but not selected) and the uncued locations (*EYE: t*(7) = −4.21; *p* = 0.012; *d* = 1.49 and HAND: *t*(7) = −3.43; *p* = 0.033; *d* = 1.21), suggesting that the central cues only modulated performance once they indicated potential motor targets. In the *EYE-HAND* condition we compared the processing rates between the motor targets and the averaged processing rates at non-targets as well as between the two effector targets. Processing rates were strongly increased at both the saccade (*t*(7) = −4.76; *p* = 0.006; *d* = 1.68) and reach target (*t*(7) = −7.40; *p* < 0.001; *d* = 2.62) compared to the non-targets. Importantly, no difference was observed between the eye and hand targets (*t*(7) = 1.74; *p* = 0.378; *d* = 0.61), indicating that attention was deployed similarly to both effector targets. We validated our approach of using the estimated processing rate parameter by additionally analyzing the observed average letter encoding (classification) probability at the different locations (i.e., the proportion of correctly reported letters for each location; Appendix **fig. A2a-b**). We observed a strikingly similar pattern of results, suggesting that the TVA parameter *v* adequately captured the deployment of attention to the different locations.

**Fig. 2.**
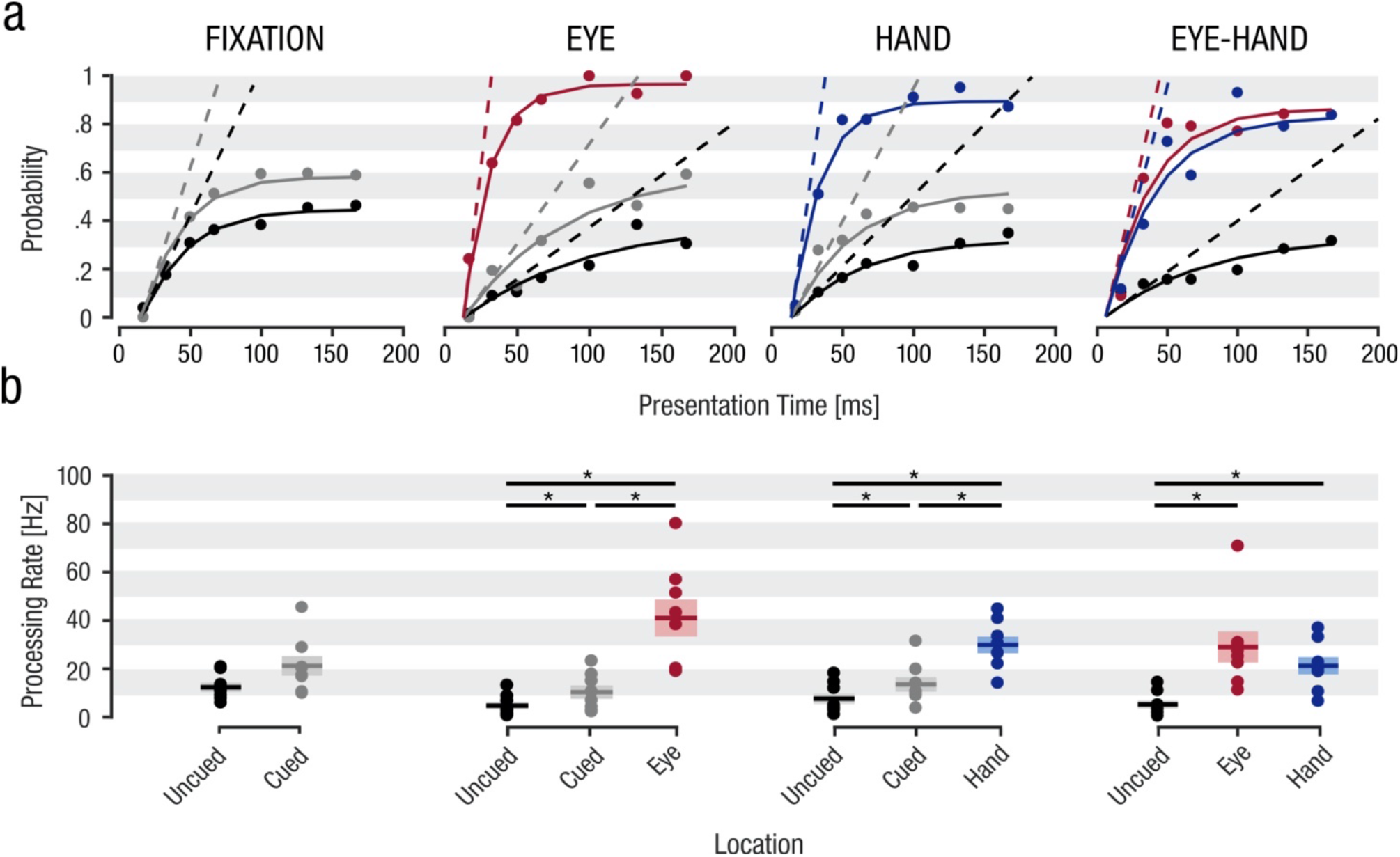
Letter encoding probabilities and processing rates at motor targets and non-targets in Experiment 1. **(a)** Letter encoding probabilities of a representative observer at the different locations as a function of letter presentation duration, separately for each experimental condition. Dots depict averaged letter encoding probabilities per target type (e.g. across all uncued locations) and presentation duration, solid lines represent the fitting curves for each target type in the different experimental conditions. Dashed lines represent the extracted processing rates (*v*). **(b)** Extracted v-parameter averaged across all participants for the respective locations and experimental conditions. Dots depict individual data points, horizontal lines represent the sample mean, and shaded areas visualize ± 1 standard error of the mean (SEM). Asterisks represent significant *p*-values < 0.05.

Noteworthy, the prioritization of item processing at the motor targets was also reflected by the order participants reported the letters: The first reported letter coincided with the letter at the saccade target in 60.4% of trials in the *EYE* condition. In the *HAND* condition the letter at the reach goal was reported first in 37.8% of trials. Similarly, in the *EYE-HAND* condition, the first reported letter was the saccade target in 51.1% and the reach target in 22.3% of trials.

Taken together, these results demonstrate that during eye and hand movement preparation, processing resources are preferably allocated towards the motor target(s): In all movement conditions the processing rate was enhanced at eye and hand targets compared to both the uncued locations as well as to cued but not selected locations. This demonstrates that the processing benefit at the motor targets is caused by the coupling between visual attention and motor target selection and cannot be explained by the mere presence of visual cues. Importantly, and in line with our hypothesis, when eye and hand movements were prepared simultaneously, we found similarly enhanced performance at both the eye and the hand target.

#### 2.2.2. Attention Capacity Across the Whole Stimulus Display

Previous studies that argued in favor of independent, effector-specific selection mechanisms showed that simultaneous eye and hand movement preparation results in parallel allocation of attention (Jonikaitis & Deubel, 2011; Hanning et al., 2018) and visual working memory benefits at their motor targets (Hanning & Deubel, 2018). Thus, the simultaneous enhancement of visual processing at two separate locations – the eye and the hand target – may be associated with an overall enhancement in attention and vSTM storage capacity over the whole visual work space; i.e., the processing capacity (*C*) of items into vSTM and the capacity of vSTM (*K*) may increase with the number of active effectors. In line with the observation that attention increases contrast sensitivity (e.g., Carrasco, 2011), we also investigated the possibility that enhanced overall attention capacity is associated with decreased perceptual thresholds (*t_0_*). An increase of processing capacity and vSTM storage capacity and a decrease of perceptual threshold would be in line with the assumption of independent, effector-specific attentional mechanisms. Alternatively, the parallel processing benefits at eye and hand targets may have concomitant costs at other, movement-irrelevant locations, such that the overall attention capacity would be independent of the number of recruited effectors. To distinguish between these two possibilities, we evaluated the overall attention capacity by modeling the item classification rate over the whole stimulus display (**fig. 1b**) and extracted the TVA parameters for each experimental condition (**fig. 3**).

**Fig. 3.**
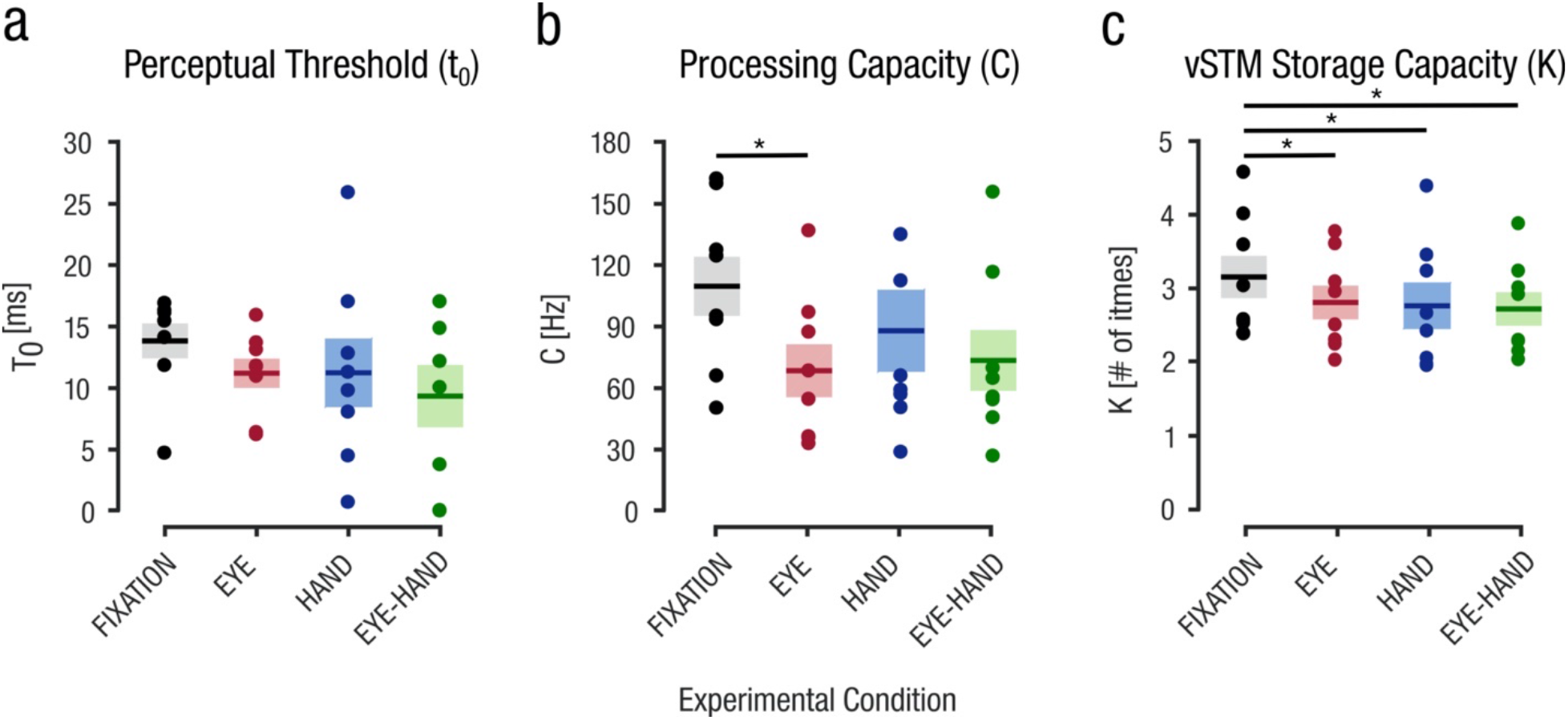
Extracted TVA whole report parameters t_0_, C, and K of Experiment 1. TVA fitting was performed for each observer and condition separately. Dots depict (**a**) individual perceptual thresholds, (**b**) processing capacities or (**c**) vSTM storage capacities. Conventions as in Figure 2.

A one-way rmANOVA with factor *experimental condition* revealed no main effect on the perceptual threshold (*F*(3,21) = 1.76; *p* = 0.186; 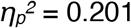) but on processing capacity (*F*(3,21) = 6.61; *p* = 0.002; 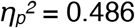) and vSTM storage capacity (*F*(3,21) = 9.92; *p* = 0.006; 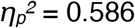). Contrary to the hypothesis of attention capacity increasing with the number of active effectors, pairwise post-hoc comparisons revealed a slight decrease for both processing capacity and vSTM storage capacity between the *FIXATION* and the movement conditions: between the *FIXATION* and *EYE* condition we observed a significant decrease of the processing capacity (*C*) (*t*(7) = 6.51; *p* = 0.002; *d* = 2.30; all other comparisons *p* > 0.077). For vSTM storage capacity, post-hoc comparisons revealed significant differences between the *FIXATION* and all movement conditions (*t*(7) = 4.09; *p* = 0.03; *d* = 1.45, *t*(7) = 6.22; *p* = 0.002; *d* = 2.20, and *t*(7) = 5.36; *p* = 0.006; *d* = 1.90, for comparisons between the *FIXATION* condition and the *EYE, HAND,* and *EYE-HAND* condition, respectively). Thus, the overall attention capacity did not rise with the number of active effectors, but rather was slightly reduced in the single and combined movement conditions compared to the *FIXATION* condition. This likely reflects dual-task costs (Künstler et al., 2018; Poth et al., 2014): Instead of focusing exclusively on the letter identification task, in the movement conditions also the arrow cues had to be interpreted and the corresponding actions had to be performed. We confirmed the results of the TVA whole report parameters *C* and *K* by additionally analyzing the average number of reported letters across presentation times (Appendix **fig. A2c**).

In sum, our results demonstrate that attention is primarily allocated towards effector targets, increasing their probability to be encoded into vSTM. When eye and hand movements are programmed simultaneously, processing resources are similarly increased at both locations and without one effector dominating the other. Given that the overall attention capacity did not increase with the number of active effectors, our results suggest that when attention is deployed at the motor targets, attentional costs should occur at the remaining, non-target locations.

## 3. Experiment 2

The second experiment was designed to assess whether the increased attention deployment at motor targets is achieved by withdrawal of processing resources from non-target locations. To give participants an incentive to deploy their attention also to non-target locations, we introduced a reward manipulation. Monetary reward has been shown to modulate voluntary selective visual attention (Della Libera & Chelazzi, 2006; for a recent review see Failing & Theeuwes, 2018), as indicated by faster reaction times and earlier N2pc components in the electroencephalogram (EEG; Kiss et al., 2009). We expected the reward manipulation in our premotor TVA paradigm to increase the attentional weights (*w_x_*) of the high-compared to the low-reward targets, reflected in higher and lower processing rates, respectively. This approach enabled us to investigate how high-reward targets compete with motor targets for attentional selection.

### 3.1. Methods

#### 3.1.1. Participants

Eight adults (age range: 25-30 years, three female) participated in this experiment (five of whom also participated in Experiment 1, including authors PK and NMH). Participants were compensated at a fixed rate of €30 but were able to additionally earn up to €30 based on their performance in the perceptual task.

#### 3.1.2. Stimuli, Apparatus, and Procedure

Stimuli, apparatus, and procedure were identical to those described for Experiment 1 with the following exceptions: (1) letters were presented either in red (20.0 cd/m^2^; 50%) or blue (20.0 cd/m^2^; 50%; **fig. 1a**). The different colors indicated whether a letter was associated with a high or low reward. The color-reward association was kept constant during the experiment but was counterbalanced across participants: they gained 100 points for each correctly reported high-reward letter, and ten points for each low-reward letter. Feedback on the points gained during a block was presented at the end of each block. At the end of the experiment, participants received an additional monetary reward according to the points gained during the entire experiment (18,481 points = €1). (2) The dynamic mask presented before and after letter presentation now consisted of red and blue lines. (3) Letter presentation durations were the same as in Experiment 1, but we added one longer presentation time (200 ms). (4) The single movement conditions of Experiment 1 showed that motor preparation modulated the distribution of attention beyond the mere presentation of a visual cue (increased performance at motor target vs. cued, not selected locations). This demonstrates that differences between our experimental conditions are explained by motor preparation rather than visual cueing. In Experiment 2 we therefore refrained from keeping the visual input constant over the conditions and only presented movement-relevant cues (i.e., no cues in the *FIXATION* condition, only one cue in the single movement conditions, and two cues in the combined movement condition).

Experimental conditions were identical to Experiment 1 (*FIXATION, EYE, HAND, EYE-HAND*). High and low-reward letters and motor targets were randomly assigned to the six possible stimulus locations with equal probability for each location. Thus, the effector(s) were equally likely to move to high or low-reward letters. The color-reward manipulation created different sub-conditions, given that each effector could either be directed to a high or a low reward target. Single effector conditions consisted of two sub-conditions: (1) effector on high-reward letter (*EYE+* or *HAND+*) or (2) effector on low-reward letter (*EYE-* or *HAND-*), and the combined movement condition consisted of four sub-conditions: (1) eye and hand on high-reward targets (*EYE+HAND+*), (2) eye on high-reward and hand on low-reward target (*EYE+HAND-*), (3) eye on low-reward and hand on high-reward target (*EYE-HAND+*), and (4) eye and hand on low-reward targets (*EYE-HAND-*).

Participants performed one training block (20 trials) of each condition, followed by 24 experimental blocks (minimum 70 trials each), split into two sessions. The block order was randomized (three *FIXATION,* six *EYE,* six *HAND,* and nine *EYE-HAND* blocks).

#### 3.1.3. Data Analysis

We applied the same trial exclusion criteria as in Experiment 1, based on which 10.22% of the trials were excluded. Data analysis and TVA modeling procedure also were identical to Experiment 1. However, instead of comparing processing rates at different locations within the experimental conditions, here we compared them across the different experimental conditions. To assess how well participants can deploy their voluntary attention according to the reward instruction when preparing single or combined movements, we calculated their ability to select high-reward letters over low-reward letters as follows:

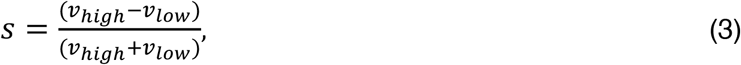

where *v_high_* and *v_low_* denote the summed processing rates of the high and low reward items, respectively. A positive reward selectivity value indicates that more high-reward letters were reported, while a negative value indicates that more low-reward items were classified. We estimated these values separately for each of the above described sub-conditions (*EYE+, EYE-, HAND+, HAND-, EYE+HAND+, EYE+HAND-, EYE-HAND+, EYE-HAND-),* to quantify how motor preparation towards or away from high reward items affects their voluntary attentional selection.

### 3.2. Results

#### 3.2.1. Allocation of Attention Across Experimental Conditions

In Experiment 1, when eye and hand movements were programmed simultaneously, we observed a similar increase of attention at both effector targets. Still, the overall attention capacity did not increase with the number of active effectors. This implies that the attention benefits at the motor targets should have concomitant costs at non-target locations. To evaluate attentional trade-offs between motor targets and non-targets, we compared the respective processing rates across the different experimental conditions (**fig 4**). We replicated our findings from Experiment 1: more processing resources were allocated to motor targets as compared to non-targets (*EYE: t*(7) = −8.55; *p* < 0.001; *d* = 3.02, *HAND: t*(7) = −4.29; *p* = 0.004; *d* = 1.52, *EYE-HAND*: *F*(2,14) = 25.01; *p* < 0.001; 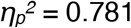). For the *EYE-HAND* condition, we again observed similarly increased processing rates at both motor targets (post-hoc comparison between eye and hand target: *t*(7) = 1.33; *p* = 0.678; *d* = 0.469). To assess a potential complementary withdrawal of attention from movement-irrelevant (non-target) locations, we ran a one-way rmANOVA with the factor *experimental condition* and observed a significant main effect (*F*(3,21) = 24.69; *p* < 0.001; 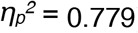). Post-hoc pairwise comparisons revealed a significant reduction in the processing rates at non-targets between the *FIXATION* condition and the *EYE* condition (*t*(7) = 5.15; *p* = 0.008; *d* = 1.82) as well as the *EYE-HAND* condition (*t*(7) = 8.68; *p* < 0.001; *d* = 3.07). We also observed a significant decrease between the *HAND* and the *EYE-HAND* condition (*t*(7) = 5.51; *p* = 0.005; *d* = 1.95) as well as between the *HAND* and the *EYE* condition (*t*(7) = −4.25; *p* = 0.023; *d* = 1.50). All other comparisons were non-significant (*p* > 0.264).

**Fig. 4.**
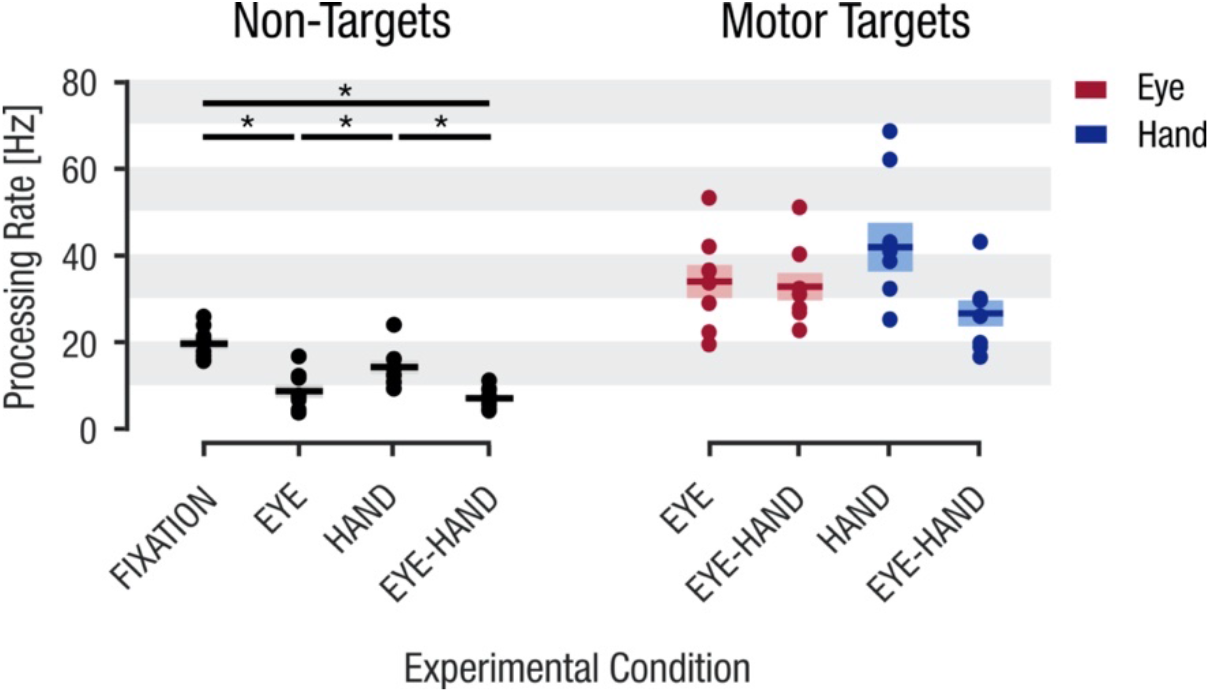
Processing rates (*v*-parameter) at non-targets and motor targets for each experimental condition in Experiment 2. Processing rates for the non-targets represent the average of the uncued locations (i.e., all six locations in the *FIXATION* condition, five in the *EYE* and *HAND* conditions, and four in the *EYE-HAND* condition). Conventions as in Figure 2.

We next assessed if there were differences in the allocation of attention between the saccade and reach targets and whether visual processing at the effector targets varied depending on whether a single or combined movement was programmed. To this end, we ran a 2 (*effector type;* eye versus hand) × 2 (*movement type;* single versus combined) rmANOVA on the estimated processing rates at the motor targets. Although there was a main effect of *movement type* (single versus combined; *F*(1,7) = 22.09; *p* < 0.002; 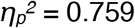), we did not find any significant differences based on the post-hoc pairwise comparisons (all *p* > 0.133). Neither the main effect of *effector type* (eye versus hand; *F*(1,7) = 0.10; *p* = 0.761; 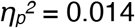), nor the interaction term (*F*(1,7) = 1.88; *p* = 0.213; 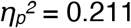) reached significance. This suggests that the shift of attention to one effector’s motor target is not hampered by movement preparation of the other effector. Instead, in order to allocate attention simultaneously to eye and hand targets, processing resources are primarily withdrawn from movement-irrelevant locations. We performed the same analysis on our data from Experiment 1 and observed a similar pattern (see Appendix **fig. A3**).

#### 3.2.2. Reward Selectivity Across Experimental Conditions

To further investigate the selective deployment of visual attention towards the effector targets at the cost of attention withdrawal from non-targets, we quantified the ability to voluntarily deploy attention while performing goal-directed movements. To this end, we biased participants’ voluntary attention by a reward manipulation: to maximize monetary reward, they should prioritize the processing of high reward letters (indicated by a specific color), and preferably name as many of these as possible. We modeled the processing rates of the high and low reward items for each sub-condition (i.e., depending on whether the movements were being prepared towards the high or low reward items). This approach also allowed us to investigate the occurrence of attentional costs *within* each experimental condition, as the comparison is not confounded by any potential differences in the overall attention capacity between experimental conditions. **Figure 5a** shows the fitted curves of a representative subject for the *EYE-HAND* condition. When both movements were directed to high-reward items (*EYE+HAND+),* processing rates were high for the high-reward items compared to low-reward items. Thus, in line with the task instructions, in this sub-condition the participant successfully allocated processing resources to maximize reward. However, already when only one effector was directed to a low-reward item (*EYE-HAND+* or *EYE+HAND-),* the ability of the participant to selectively report high over low-reward items decreased remarkably. Crucially, when both effectors targeted low-reward items (*EYE-HAND-*), the participant even preferably reported low-reward items, demonstrating that motor preparation overrode voluntarily attempts to attend elsewhere.

**Fig. 5.**
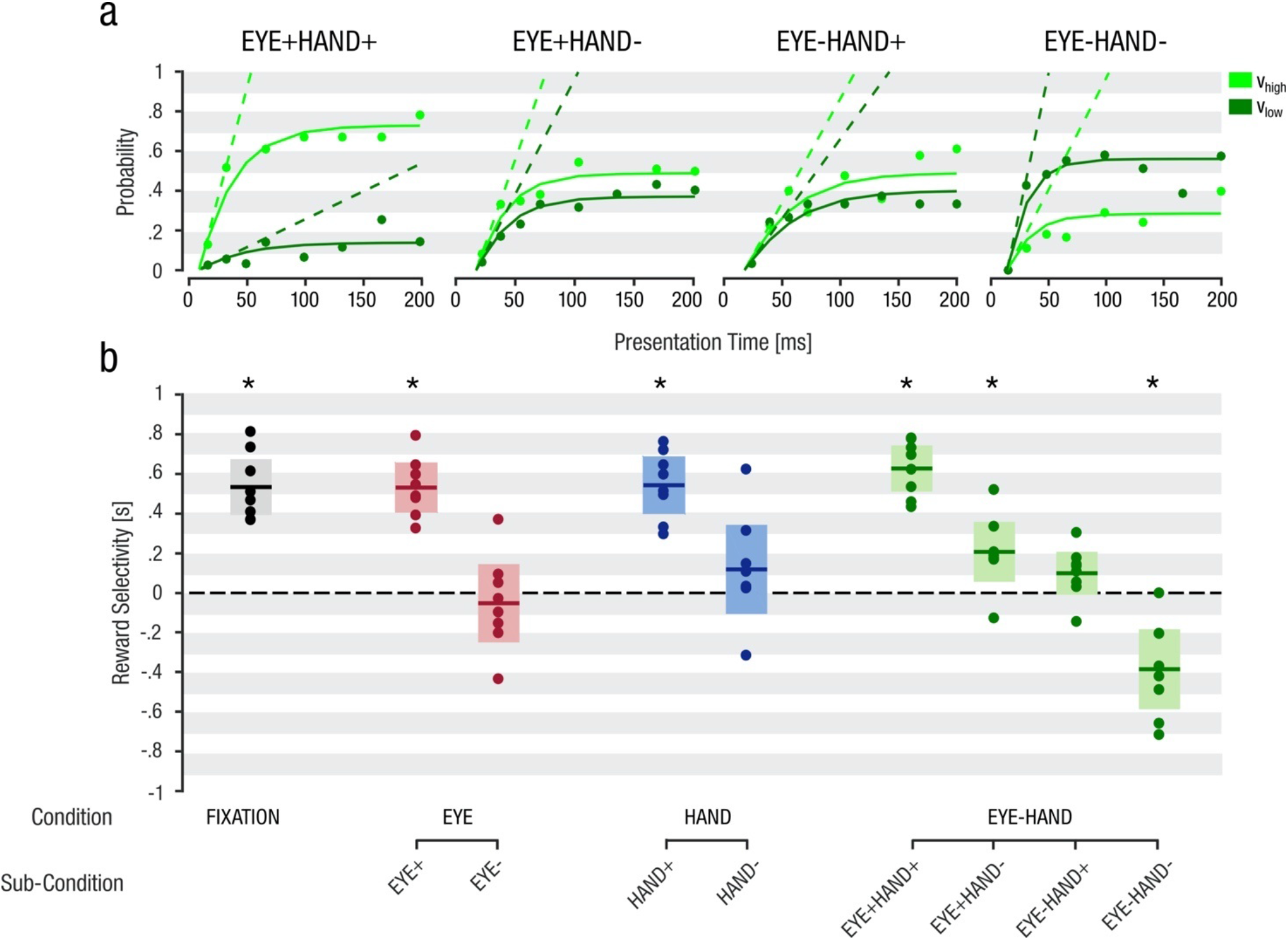
**(a)** Average fitting curves of a representative participant for high reward items (light green) and low reward items (dark green) in the *EYE-HAND* condition. Dashed lines represent the respective averaged processing rate. **(b)** Average reward selectivity (defined as the relative difference between high- and low-reward encoding rates) across all participants. Data was modeled for each sub-condition and participants separately. Dots represent individual subject data points. Solid lines depict the sample mean and shaded areas show 95% CI.

The extracted measurement of reward selectivity (*s*) shows that all participants exhibited a strikingly similar pattern of results (right panel of **fig. 5b**). We calculated the mean and 95% confidence intervals across participants for each sub-condition to assess their difference from 0 (i.e., no selective encoding preference for neither high-nor low-reward items; **fig. 5b**). As expected, in the *FIXATION* condition participants allocated significantly more processing resources towards high-reward items than low-reward items (*mean* = 0.53; 95% CI [0.39, 0.67]), demonstrating that our reward manipulation indeed biased their voluntary allocation of attention. During the preparation of goal-directed actions, item selection depended primarily on the location of the motor targets: participants only continued reporting more high than low-reward items as long as the motor target(s) were congruent with the high-reward item locations (*EYE+: mean* = 0.53; 95% CI [0.41, 0.65], *HAND+: mean* = 0.54; 95% CI [0.40, 0.68], and *EYE+HAND+: mean* = 0.62; 95% CI [0.51, 0.74]). However, when one movement was programmed towards a low-reward item, participants selected this item, which declined their ability to prioritize high-reward over low-reward items. This was true for the single movement conditions (*EYE-: mean* = −0.05; 95% CI [−0.25, 0.15] and *HAND-: mean* = 0.12; 95% CI [−0.11, 0.34]) and the *EYE-HAND* condition (*EYE+HAND-: mean* = 0.21; 95% CI [0.06, 0.36] and *EYE-HAND+: mean* = 0.10; 95% CI [−0.01, 0.21]). Critically, when both effectors targeted low-reward items in the *EYE-HAND* condition, the allocation of attentional resources was aligned with the motor programs (i.e., more processing resources were allocated towards low-reward items; *EYE-HAND-: mean* = −0.38; 95% CI [−0.58, −0.18]).

This demonstrates that even when participants aim to voluntarily attend to high-reward but non-motor targets, attention nonetheless is withdrawn from the voluntarily selected items and shifts towards the motor targets. Our limited processing resources are thus primarily allocated towards motor targets, leaving only few resources to be voluntarily deployed to movement-irrelevant locations.

## 4. Discussion

In our novel premotor TVA paradigm, participants reported briefly presented letters while preparing eye, hand, or simultaneous eye-hand movement. To the best of our knowledge, this study is the first to apply TVA to investigate attentional dynamics during action preparation. By simultaneously assessing visual processing at motor targets and movement-irrelevant locations, we show that TVA is a sensitive and innovative tool to evaluate premotor attentional dynamics.

In TVA, objects in the visual work space compete for successful encoding into visual short-term memory (vSTM). Attentional weights assigned to each object determine the winner(s) of this race. The neurocomputational account of TVA (NTVA; Bundesen et al., 2005; 2011) proposes a two-stage process of distributing attentional weights. Initially, processing resources are distributed randomly across all objects. At the end of a first perceptual cycle, attentional weights are computed for each object based on their momentary importance and stored in a priority map. During a second perceptual cycle, processing resources are selectively allocated across the objects according to their attentional weights. Based on our data, we propose that motor preparation strongly influences the distribution of these attentional weights: To successfully perform movements, attentional weighting is increased at motor targets, while being decreased at movement-irrelevant locations. Such a close link between attentional weighting and motor target selection was previously supported by a TVA-based computational model of visual attention that derived priority maps from attentional weights to determine saccade targets (Wischnewski et al., 2010).

Whereas earlier reports showed that TVA can be extended to account for dual task situations (Logan & Duncan, 2001), including motor-cognitive dual task-costs (Künstler et al., 2018), our data show – for the first time – how movement preparation directly influences the selection of competing objects in the visual work space. Our study not only demonstrates that TVA can accommodate attentional mechanisms in relation to selection-for-action (Allport, 1987; Schneider, 1995), but also incorporates premotor attention in the broader context of the biased competition framework (Desimone & Duncan, 1995), as previously suggested by Smith and Schenk (2012). Rather than describing attention as a moving spotlight, this framework proposes that attentional mechanisms resolve the competition of objects in the visual work space for a limited processing capacity and control of behavior.

Our novel approach allowed us to shed light on the attentional dynamics underlying motor target selection during simultaneous eye-hand movements. The following section discusses our results with respect to previous work and concludes with an updated framework of attentional dynamics underlying simultaneous action preparation (**fig. 6**). In line with earlier studies (Hanning et al., 2018; Jonikaitis & Deubel, 2011), we found that visual attention was equally and in parallel allocated to both effector targets during simultaneous eye-hand movement preparation. This is in contrast to studies reporting that the eye carries more attentional weight than the hand during simultaneous eye hand movements (Khan et al., 2011; Stewart et al., 2019; see **fig. 6a** for predictions based on *eye-dominance*). Instead, our results show that attention was similarly increased at both the eye and the hand target, without one effector dominating over the other. Moreover, the deployment of attention to one effector target was not hampered by concurrent motor preparation of the other effector system (note that small performance reductions at the motor targets between the single and combined movement conditions can be explained by an overall increase in task difficulty when performing two instead of one movement. The finding of separate attentional benefits at eye and hand movement targets has been interpreted as evidence for independent, effector-specific attentional mechanisms (Hanning et al., 2018; Jonikaitis & Deubel, 2011; see **fig. 6b** for predictions based on *independent systems*). This pattern, however, may likewise be achieved by attentional withdrawal from movement-irrelevant locations – with the overall attention capacity being unaffected by the number of active effectors. Going beyond traditional premotor paradigms measuring visual attention only at one location at a time (Deubel et al., 1998; Hanning et al., 2018; Hanning et al., 2019; Jonikaitis & Deubel, 2011; Khan et al., 2011), our premotor TVA approach allows us to test this alternative explanation by assessing premotor visual processing across the entire visual work space, including motor targets as well as movement-irrelevant locations. Findings reveal that processing resources are primarily allocated towards motor targets, with separate benefits at both targets before simultaneous eye-hand movements. Importantly, however, these motor target benefits were associated with attentional costs arising at non-target locations, yielding a constant attentional capacity that does not increase with the number of active effectors. Such withdrawal of processing resources from movement irrelevant locations is in line with a recent study showing attentional suppression at non-saccade targets (Khan et al., 2015).

**Fig. 6.**
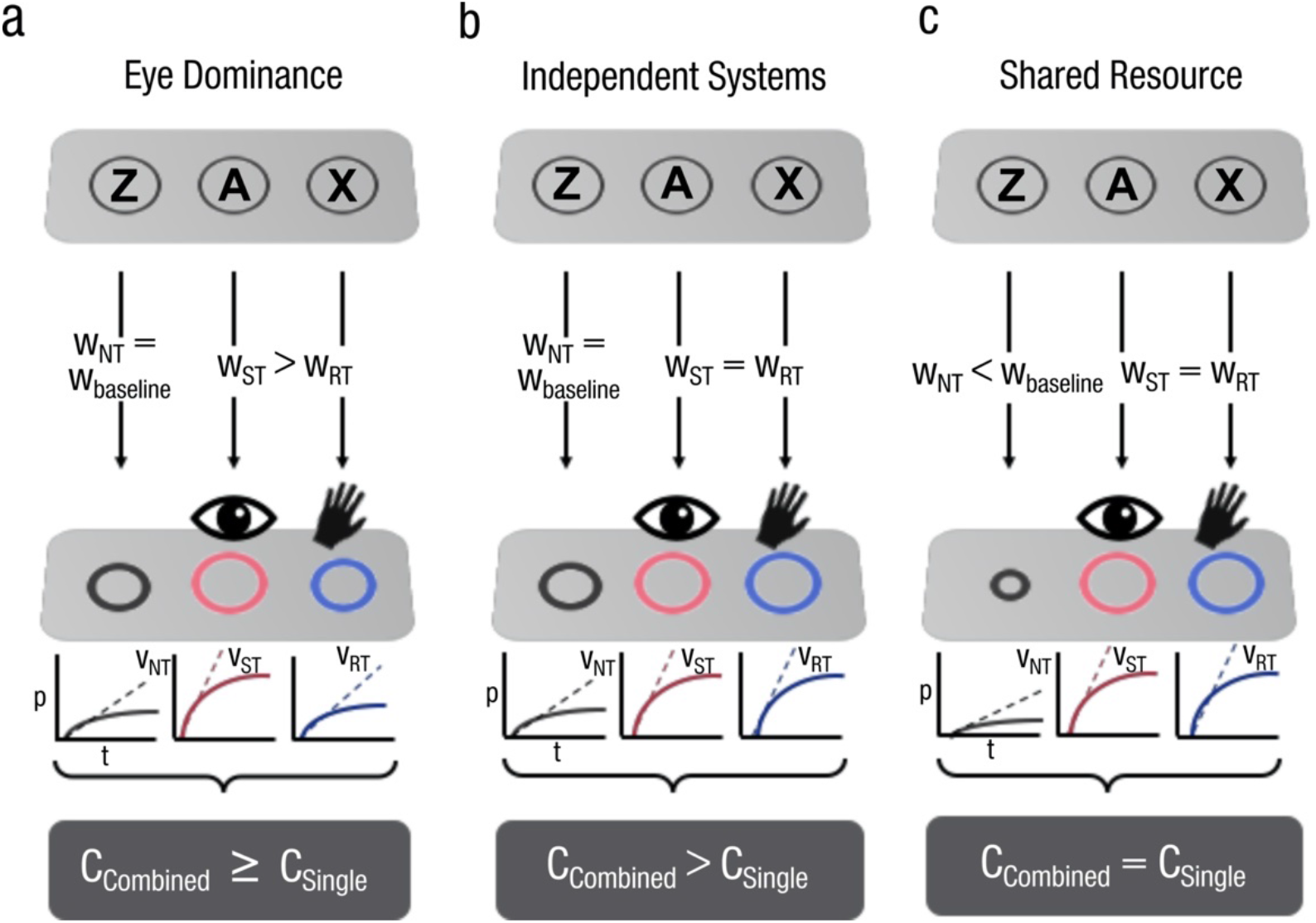
Current models of attentional dynamics underlying simultaneous eye-hand movement preparation. **(a)** The *Eye Dominance* model proposes that the eye predominantly guides attention during the planning of simultaneous eye-hand movements. In TVA, this would lead to higher attentional weights assigned to the saccade target than the reach target. The overall attentional capacity will only slightly, if at all, increase with the number of active effectors. **(b)** The *Independent Systems* model assumes similarly increased attentional weighting at the saccade and reach targets. Importantly, this model proposes that there is no attentional competition between the two effectors (i.e., the attentional deployment to one effector target is not hampered by the other effector). If attentional benefits at eye and hand targets in fact are independent of each other, the overall attention capacity should be higher in the combined (*EYE-HAND*) compared to the single (*EYE* or *HAND*) movement condition. Note that neither the eye dominance nor the independent systems model makes predictions regarding attentional costs at non-motor targets. **(c)** In contrast, our *Shared Resource* model proposes that attention is withdrawn from movement irrelevant locations in order to achieve equal attentional deployment at eye and hand targets. Assuming that a single, shared attentional resource is dynamically deployed over the visual work space, the overall attention capacity should be fixed and unaffected by the number of active effectors (i.e., *C* is similar in the single and combined movement conditions). NT = non-target, ST = saccade target, RT = reach target, w = attentional weight, v = processing rate (initial rise in object encoding probability p with increasing presentation time t), C = processing capacity. Differently sized circles represent different attentional weighting.

To further investigate the withdrawal of processing resources from non-target locations, we introduced a reward manipulation in our second experiment. Similar to increased attentional weighting of motor targets, high reward targets are prioritized for attentional selection (Failing & Theeuwes, 2018), which is reflected in shorter reaction times and enhanced N2pc components (Kiss et al., 2009). Thus, combining our premotor TVA paradigm with a reward manipulation allowed us to assess how premotor attention interacts with voluntarily deployed attention to maximize reward. As expected, we observed increased processing at high-reward compared to low-reward object locations when no motor action was required. This reward selectivity, however, sharply decreased once an eye or hand movement was planned away from the voluntarily attended objects, and completely vanished when both effectors were directed away from high-reward objects. This demonstrates that processing resources are withdrawn from movement-irrelevant locations in order to shift attention to motor targets, even when participants aim to select visual content at non-target locations. Crucially, while motor preparation resulted in attention withdrawal from non-target locations, we did not find pronounced attentional competition between the eye and hand (i.e., neither of the two movements affected the benefit at the other effector target).

Based on our findings we propose an updated framework of attentional mechanisms underlying simultaneous eye-hand movements. Our finding of equal attentional weighting of eye and hand targets argues against *eye-dominance* (**fig. 6a**). In line with *independent attentional systems* (**fig. 6b**), our data show a parallel attention allocation without pronounced competition between eye and hand targets. However, revising the independent systems interpretation, we moreover demonstrate that this simultaneous enhancement of processing resources at different effector targets is associated with attentional costs at non-motor targets. Our observation that attentional premotor target benefits are accompanied by non-motor costs suggests that the overall attention capacity remains fixed. In fact, our TVA analysis reveals that the parameter *C* (processing capacity) did not increase with the number of active effectors. Our results are thus best explained by a *shared resource* model (**fig. 6c**), in which a fixed processing resource is flexibly distributed across the visual work space according to momentary behavioral relevance. A parallel allocation of visual attention to motor targets of different effectors without competition (Hanning et al., 2018; Jonikaitis & Deubel, 2011) may still be possible as long as sufficient processing resources can be withdrawn from movement-irrelevant locations. Noteworthy, previous studies investigating premotor attention during eye-hand movements (Hanning et al., 2018; Jonikaitis & Deubel, 2011; Khan et al., 2011) were not able to analyze the distribution of attention over multiple items.

Using alternative-forced-choice discrimination tasks, where participants typically perform close to chance at non-targets, these paradigms were neither designed to precisely measure the allocation of attention at movement-irrelevant locations, nor to assess the overall attention capacity. As our results show, the analysis of attention distribution over the whole visual work space sheds further light onto the mechanisms underlying parallel attention allocation at multiple effector targets. Thus, our premotor TVA paradigm goes beyond traditional sensitivity measures and provides a powerful tool for a more thorough analyses of premotor visuospatial attention across the visual scene.

## 5. Conclusion

TVA-based assessments of visual attention have been widely applied in both clinical (Habekost, 2015) and basic research (Bundesen & Habekost, 2014) – but solely under fixation conditions. By combining TVA with motor tasks, we are the first to show that this framework can also be used to evaluate the dynamics of visual attention associated with motor preparation. Our study demonstrates that the commonly observed coupling of attention and goal-directed actions can be accommodated within the broader framework of biased competition (Desimone & Duncan, 1995; Smith & Schenk, 2012) and provides several new insights into the attentional mechanism underlying the preparation of combined eye-hand movements.

First, while eye and hand movements draw an equal amount of attentional resources towards their motor targets, allegedly not interacting with each other, we show that attentional costs occur at non-motor targets in order to allocate processing resources to the movement goals. Second, our reward manipulation demonstrates that these attentional costs cannot be overcome even when participants have a strong incentive to attend to non-motor targets. Third, since the overall attention capacity did not increase with the number of active effectors, our findings reveal that eye and hand movements share a common attentional resource.

In summary, the remarkable feature of TVA – the ability to quantify both, attention deployment to individual objects as well as overall attention capacity – allows to thoroughly analyze the allocation of attention in action contexts, overcoming limitations of previous studies that could only measure perceptual sensitivity at a single stimulus location at a time. Future research employing single-unit neurophysiology, neuroimaging, and computational techniques is required to advance our understanding of the neural substrates underlying a shared attentional resource to which both the eye and the hand movement system have simultaneous and equal access.

## Author Note

The data is available on the Open Science Framework (https://osf.io/h9342/). No part of the study was pre-registered prior to the research being conducted.

## Conflict of interest

The authors declare no conflicts of interest.

## Author Contributions

All authors developed the study concept and contributed to the study design. P.K. and N.M.H. collected and analyzed the Data. All authors interpreted the data. P.K. drafted the manuscript, and H.D. and N.M.H. provided critical revisions. All authors approved the final version of the manuscript for submission.

## Acknowledgements

The authors thank Marleen Haupt, Natan Napiórkowski, Christian Poth, and members of the Deubel laboratory for helpful advices and discussions. This work was supported by the Deutsche Forschungsgemeinschaft (DFG) [DE336/5-1].

## Appendices

**Fig. A1.**
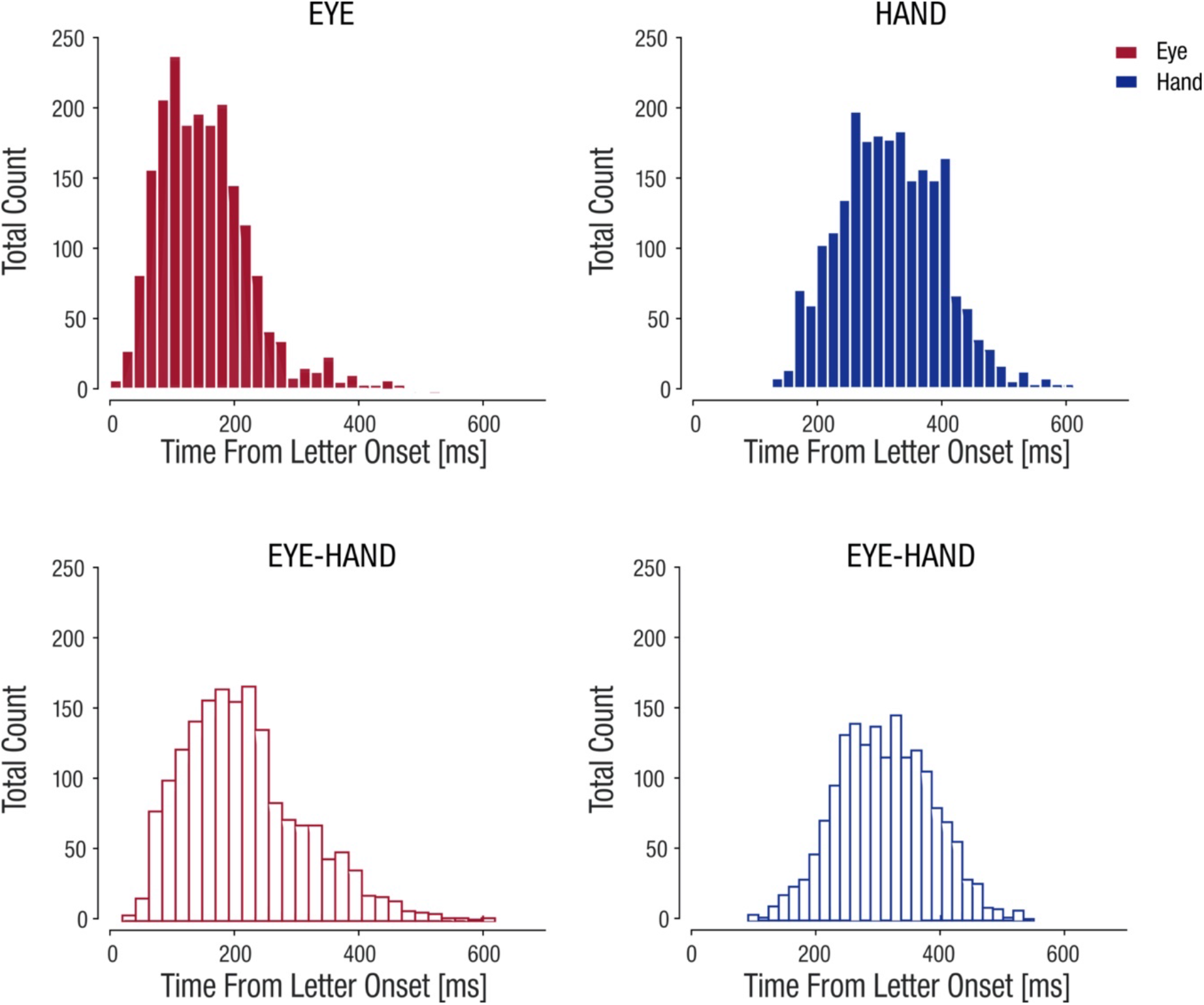
Distribution of movement latencies locked to letter probe onset in the different movement conditions. Neither eye (red) nor the hand (blue) movement latencies were affected (inhibited) by letter probe onset – which would be evident in a frequency depression at approximately 90 to 100 ms (Reingold & Stampe, 2002). This was true for both the single (solid bars) and combined (open bars) movement conditions.

**Fig. A2.**
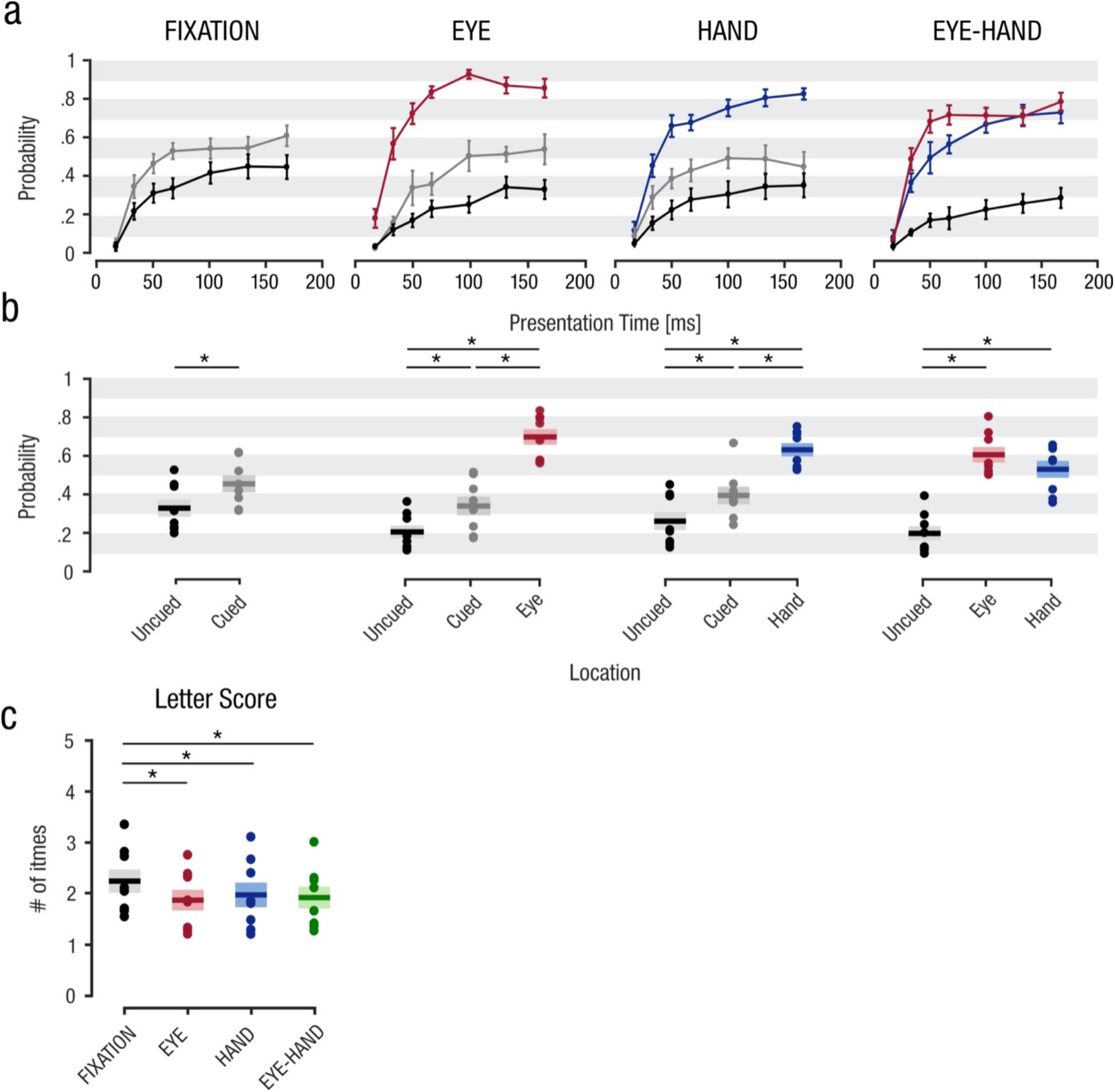
Observed letter report performance in Experiment 1. (a) Mean letter encoding probability at the different locations within experimental conditions plotted as a function of presentation times. (b) Letter report performance averaged across presentation times. In accordance with the analyses of the estimated processing rates (*v;* **fig. 4**), increased deployment of attention is observed at the motor targets in both single movement conditions and in the combined movement condition. While attention at cued but non-selected locations was increased compared to uncued locations, significantly more attention was allocated to the motor targets. Note that in the combined movement condition, letter encoding probability was >0.5 for the longer presentation times at the eye and the hand target, indicating that attention was deployed in parallel to both effector targets. (c) Total number of correctly reported letters averaged across presentation times for each experimental condition. Similar to the TVA parameter-based assessment of attention (*C*) and vSTM capacity (*K),* overall performance did not increase with the number of active effectors. Instead, small decreases in performance between the fixation and movement conditions are observed. Asterisks indicate significant post-hoc pairwise comparisons (*p* < 0.05). Errorbars denote ±1 SEM. Other conventions as in Figure 2.

**Fig. A3.**
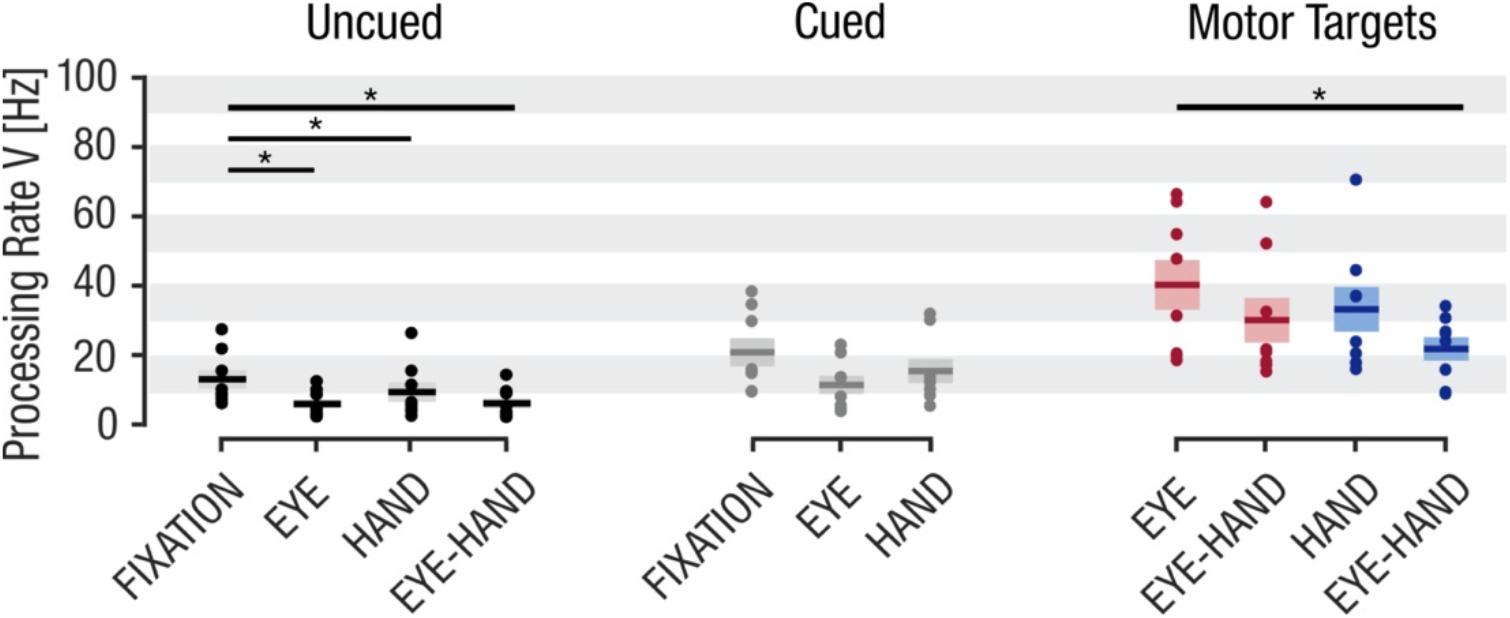
Comparison of processing rates (*v*) at uncued, cued but non-selected, and motor targets across conditions of Experiment 1. A significant withdrawal of attention was observed at the uncued locations in the movement conditions compared to the fixation condition. There was no significant decrease between the single (*EYE* or *HAND*) and combined (*EYE-HAND*) movement conditions. This may be related to a floor effect, as processing rates were already low in the single movement conditions in order to shift attention to the motor targets. A similar (non-significant) trend was also evident at the cued, but non-selected locations. Attention was increased similarly at the eye and hand targets, irrespective of whether a single or combined movement was planned. Asterisks indicate significant post-hoc pairwise comparisons (*p* < 0.05). Conventions as in Figure 2.

## Notes

### Competing Interest Statement

The authors have declared no competing interest.

